# Neural patterns reveal single-trial information on absolute pitch and relative pitch perception

**DOI:** 10.1101/672675

**Authors:** Simon Leipold, Marielle Greber, Silvano Sele, Lutz Jäncke

## Abstract

Pitch is a fundamental attribute of sounds and yet is not perceived equally by all humans. Absolute pitch (AP) musicians perceive, recognize, and name pitches in absolute terms, whereas relative pitch (RP) musicians, representing the large majority of musicians, perceive pitches in relation to other pitches. In this study, we used electroencephalography (EEG) to investigate the neural representations underlying tone listening and tone labeling in a large sample of musicians (n = 105). Participants performed a pitch processing task with a listening and a labeling condition during EEG acquisition. Using a brain-decoding framework, we tested a prediction derived from both theoretical and empirical accounts of AP, namely that the representational similarity of listening and labeling is higher in AP musicians than in RP musicians. Consistent with the prediction, time-resolved single-trial EEG decoding revealed a higher representational similarity in AP musicians during late stages of pitch perception. Time-frequency-resolved EEG decoding further showed that the higher representational similarity was present in oscillations in the theta and beta frequency bands. Supplemental univariate analyses were less sensitive in detecting subtle group differences in the frequency domain. Taken together, the results suggest differences between AP and RP musicians in late pitch processing stages associated with cognition, rather than in early processing stages associated with perception.

## Introduction

Music, speech, and many environmental sounds have acoustic waveforms that repeat over time. The repetition rate of these sounds is perceived as pitch (Plack et al., 2005). Pitch is not perceived equally by all humans: Patients with congenital amusia (tone deafness) are limited in judging whether a pitch is higher or lower than a previous pitch (Peretz, 2016). Healthy participants show a consistent preference to perceive either the fundamental pitch or the spectral pitch of a sound (Schneider et al., 2005). And then, there is a special group of musicians which effortlessly perceive, recognize, and name pitches in absolute terms and without the help of a reference pitch (Deutsch, 2013). These absolute pitch (AP) musicians stand in striking contrast to the large majority of relative pitch (RP) musicians which perceive pitches in relation to other pitches (McDermott and Oxenham, 2008; Miyazaki and Rakowski, 2002).

The neural correlates of AP and RP have been extensively studied using electroencephalography (EEG), yielding various findings regarding the auditory event-related potential (ERP) (Behroozmand et al., 2014; Bischoff Renninger et al., 2003; Burkhard et al., 2019; Crummer et al., 1994; Elmer et al., 2013; Greber et al., 2018; Hantz et al., 1992; Itoh et al., 2005; Klein et al., 1984; Leipold et al., 2019b; Rogenmoser et al., 2015; Tervaniemi et al., 1993; Wayman et al., 1992). All of these studies exclusively applied univariate analysis, which includes a preselection of electrodes and time windows to be analyzed. The classical univariate approach only utilizes a small portion of the collected data and disregards that the EEG signal is inherently multivariate (Michel and Murray, 2012). Furthermore, these studies calculated grand averages across numerous trials (Luck, 2014) and thus disregarded within-participant variance, which demonstrably contains information on perception and cognition (Ratcliff et al., 2009).

Recent methodological developments within the framework of “brain decoding” have enabled the successful extraction of information from single trials in spite of their low signal-to-noise ratio (Blankertz et al., 2011; Grootswagers et al., 2016). An increasing number of studies have used a decoding framework to analyze noninvasively obtainable electrophysiological signals (Cichy et al., 2014; Crouzet et al., 2015; King et al., 2013; Sankaran et al., 2018; Schaefer et al., 2011). In the context of EEG decoding, it is useful to characterize the EEG signal as a time series of multivariate topographical patterns dynamically changing over time (Michel and Koenig, 2018). When transformed to the frequency domain, these topographical patterns become frequency-specific. The patterns can be conceptualized as neural representations underlying the perceptual and cognitive processing that is performed in the context of experimental conditions. In other words, the patterns contain information about the experimental conditions, which can be decoded using multivariate pattern analysis (Grootswagers et al., 2016; Haxby et al., 2014). To fully exploit the excellent temporal resolution of EEG, multivariate pattern analysis is best applied to separate time samples of the signal (Hausfeld et al., 2012). For each participant, a machine learning classifier is trained and tested (e.g., using cross-validation) at each time sample to differentiate between the experimental conditions (e.g., using the data from single trials). The success of this classification is quantified, for example, by the decoding accuracy (i.e. the fraction of trials that are correctly classified). This decoding accuracy in turn quantifies the representational (dis)similarity of the neural representations underlying the perceptual and cognitive processing in the different experimental conditions; the more similar the neural representations, the lower the decoding accuracy (Grootswagers et al., 2016; Kriegeskorte et al., 2008).

In this study, we examined the neural representations underlying pitch perception in AP and RP musicians using time-resolved and time-frequency-resolved single-trial EEG decoding. Participants performed a pitch-processing task with two experimental conditions: a listening condition and a labeling condition. Cognitive models of AP suggest that pitch labeling is the most crucial component distinguishing AP from RP musicians (Levitin, 1994; Levitin and Rogers, 2005). More specifically, the label is part of the cognitive representation of the pitch and is automatically activated when a particular pitch is perceived (Levitin and Rogers, 2005). This automaticity has been empirically demonstrated using Stroop-like tasks, in which AP musicians responded slower in trials where the pitch of a tone did not match a simultaneously presented label (Akiva-Kabiri and Henik, 2012; Hsieh and Saberi, 2008; Itoh et al., 2005; Schulze et al., 2013). Thus, we predicted that the neural representations underlying listening and labeling show high similarity in AP musicians because they automatically label pitches during listening and, to comply with the task demands, also label pitches during labeling. On the contrary, RP musicians do not label tones during listening, but they must use their RP ability during labeling to follow the instructions of the task. The labeling of pitches using RP involves additional cognitive processes not present during simple listening (Itoh et al., 2005; Zatorre et al., 1998). Thus, we predicted a lower representational similarity of listening and labeling in RP musicians. We tested these predictions using the EEG data of a large sample of musicians (n = 105) by comparing time series of representational similarity between AP musicians (n = 54) and RP musicians (n = 51). We also performed classical univariate analyses to maintain comparability with previous studies. Finally, we employed a behavioral Stroop-like task to assess whether the postulated labeling automaticity is also present in our sample.

## Materials and methods

### Participants

We analyzed the data of 105 participants, who were recruited in the context of a large project investigating AP (Brauchli et al., 2019; Burkhard et al., 2019; Greber et al., 2018; Leipold et al., 2019a). The EEG data of 104 of these participants was previously analyzed as part of a direct replication study using a different group assignment and methodology (Leipold et al., 2019b). All participants were either professional musicians, music students, or highly trained amateurs, aged between 18 and 44 years. The participants’ assignment to the groups of AP musicians or RP musicians was based on self-report. None of the participants reported any neurological, audiological, or severe psychiatric disorders. We confirmed the absence of hearing loss in all participants using pure tone audiometry (ST20, MAICO Diagnostics, Berlin, Germany). The demographical data (sex, age, handedness) and part of the behavioral data (tone-naming proficiency, musical aptitude, and musical experience) were collected using an online survey tool (http://www.limesurvey.org/). We used a German translation of the Annett questionnaire to verify the self-reported handedness (Annett, 1970). Musical aptitude was assessed using the Advanced Measures of Music Audiation (AMMA) (Gordon, 1989). To probe the RP ability of the participants, we employed a musical interval identification test. Crystallized intelligence was estimated in the laboratory using the Mehrfachwahl-Wortschatz-Intelligenztest (MWT-B) (Lehrl, 2005), and fluid intelligence was estimated using the Kurztest für allgemeine Basisgrößen der Informationsverarbeitung (KAI) (Lehrl et al., 1991). The participants provided written informed consent and were paid for their participation. The study was approved by the local ethics committee (http://www.kek.zh.ch/) and conducted according to the principles defined in the Declaration of Helsinki.

### Tone-naming test

We employed a tone-naming test to assess the tone-naming proficiency of the participants (Oechslin et al., 2010). Participants had to name both the chroma and the octave (e.g., E4) of 108 pure tones presented in a pseudorandomized order. The test included all tones from C3 to B5 (twelve-tone equal temperament tuning, A4 = 440 Hz). Each of the tones was presented three times. The tones had a duration of 500 ms and were masked with 2,000 ms Brownian noise presented immediately before and after each tone. The tone-naming score was calculated as the percentage of correct chroma identifications without considering octave errors (Deutsch, 2013); the chance level was at 8.3%.

### Stroop-like task

To empirically establish the labeling automaticity in AP musicians, we applied a behavioral audio-visual Stroop-like task (Akiva-Kabiri and Henik, 2012; Hsieh and Saberi, 2008; Itoh et al., 2005; Schulze et al., 2013; Stroop, 1935). The auditory stimuli consisted of five pure tones (duration = 500 ms, 10 ms linear fade-in; 50 ms linear fade-out), which were created using Audacity (version 2.1.2, http://www.audacityteam.org/). The tones corresponded to C4 (262 Hz), D4 (294 Hz), E4 (330 Hz), F4 (349 Hz), and G4 (392 Hz) in twelve-tone equal temperament tuning. The visual stimuli were comprised of the musical notations (quarter notes in treble clef) of the same tones. Participants simultaneously heard a sine tone and viewed a musical note on a computer screen. They were asked to identify the visually presented musical notes as fast and as accurately as possible by button press (C, D, E, F, or G) while ignoring the tones. The label of the tone and the name of the musical note were either congruent or incongruent. If AP musicians are unable to suppress labeling, they are expected to experience more cognitive interference in incongruent trials than RP musicians. This would be reflected by larger response time differences between congruent and incongruent trials in AP musicians.

### Experimental procedure

All stimuli, the stimulus presentation scripts, and the raw data are available online (https://dx.doi.org/10.17605/OSF.IO/7QXJS).

#### EEG experiment

During EEG data acquisition, the participants performed a pitch processing task with a listening and a labeling condition (Itoh et al., 2005). The stimuli encompassed three pure tones (262 Hz, 294 Hz, and 330 Hz), corresponding to C4, D4, and E4, which were created using Audacity. The tones had a duration of 350 ms (10 ms linear fade-in; 50 ms linear fade-out) and were presented at a sound pressure level of 75 dB via on-ear headphones (HD 25-1, Sennheiser, Wedemark, Germany) using Presentation software (version 18.1, www.neurobs.com). Both experimental conditions consisted of 180 trials presented in a randomized order; each of the pure tones was presented 60 times. In each trial, first, a pure tone was presented. This tone was followed by a jittered inter-stimulus interval (duration = 900–1,100 ms). Then, an auditory cue (10 ms of pink noise; linear fade-in = 2 ms, linear fade-out = 2 ms) was presented to indicate to the participants that they should respond by a key press. After the cue, there was a silent period until a response was given, followed by an inter-trial interval of 1,000 ms. During the task, a screen in front of the participants showed a black fixation cross on a gray background. The conditions only differed in the instructions given to the participants: In the listening condition, the participants were instructed to listen to the tones and press a neutrally marked key in response to the auditory cue, irrespective of the chroma of the tones. In the labeling condition, they were instructed to label the tones by pressing one of three corresponding keys marked with the tone names (C, D, and E). By instructing to respond only after the cue, we avoided a contamination of the EEG by motor artifacts. In both conditions, the participants were also instructed to respond as quickly and as accurately as possible and not to respond verbally. The order of the conditions was fixed across participants, with the listening condition always preceding the labeling condition. The whole task had a duration of 20 minutes.

#### EEG data acquisition and preprocessing

The EEG was continuously recorded using an electrode cap (Easycap, Herrsching, Germany) with 32 Ag / AgCl electrodes arranged according to an extended 10/20 system (Fp1, Fp2, F7, F3, Fz, F4, F8, FT7, FC3, FCz, FC4, FT8, T7, C3, Cz, C4, T8, TP9, TP7, CP3, CPz, CP4, TP8, TP10, P7, P3, Pz, P4, P8, O1, Oz, O2) in combination with a BrainAmp amplifier (Brain Products, Munich, Germany). The sampling rate was 1,000 Hz, the data was online bandpass-filtered between 0.1 Hz and 100 Hz, the reference electrode was placed on the tip of the nose, and electrode impedance was kept below 10 kΩ throughout data acquisition by applying an electrically conductive gel.

The preprocessing of the EEG data was performed in BrainVision Analyzer (Version 2.1, https://www.brainproducts.com/). We used slightly different settings for the time-resolved EEG decoding and the time-frequency-resolved EEG decoding. In the preprocessing for the time-resolved EEG decoding, first, we bandpass-filtered the data from 0.5 Hz to 20 Hz (48 dB/octave) and applied a notch filter of 50 Hz. Next, we corrected vertical and horizontal eye movement artifacts using independent component analysis (Jung et al., 2000). Then, we removed the remaining artifacts using an automatic raw data inspection (removal criteria: amplitude gradient > 50 μV/ms, amplitude difference > 100 μV, amplitude minimum/maximum > −100 μV / +100 μV). Finally, we segmented the continuous data into epochs of 900 ms (−100 ms to +800 ms relative to stimulus onset) and applied a baseline correction using the time interval between −100 and 0 ms relative to stimulus onset. In the preprocessing for the time-frequency-resolved EEG decoding, we applied a high-pass filter of 0.5 Hz (48 dB/octave) and a notch filter of 50 Hz, but we did not use a low-pass filter (Cohen, 2014). The artifact correction procedure was identical to the time-domain analysis, but we used more liberal removal criteria during the automatic raw data inspection (amplitude gradient > 100 μV/ms, amplitude difference > 200 μV, amplitude minimum/maximum > −200 μV / +200 μV). Lastly, we segmented the data into epochs of 2,500 ms (−1,000 ms to +1,500 ms relative to stimulus onset) and normalized the time-frequency power values (dB transformation) using the time interval between −500 and −100 relative to stimulus onset as a baseline. The prolonged epoch size avoided a contamination of the time-frequency representation by edge artifacts (Cohen, 2014).

### Statistical analysis of the behavioral data

The statistical analysis of the behavioral data was performed in R (version 3.3.2, https://www.r-project.org/). The participant characteristics were compared between the groups using Welch’s t-tests. For the Stroop-like task, we analyzed the response times of the correct trials; trials with a response time shorter or longer than 2 standard deviations of the participant-and-condition-specific mean response time were excluded from the analysis. To calculate the size of the Stroop-effect for each participant, we subtracted the response times of the congruent trials from the response times of the incongruent trials. The differences in response times (incongruent minus congruent) were then subjected to a group comparison using Welch’s t-tests. Effect sizes in the context of t-tests are given using Cohen’s *d*. The significance level was set to α = 0.05 for all behavioral analyses.

### Time-resolved single-trial EEG decoding

The goal of the EEG decoding analyses was to investigate the similarity of the dynamic neural representations underlying tone listening and tone labeling in AP musicians compared to RP musicians. As part of the time-resolved single-trial EEG decoding, we trained a machine learning classifier to differentiate between the experimental conditions at each time sample of the time-domain EEG signal. As a result, we obtained a dynamic measure of representational (dis)similarity operationalized by the time series of decoding accuracies per participant. The decoding accuracy represents the information available in an EEG topography at a given time sample, in our case, about the differences of neural representations underlying the experimental conditions (Grootswagers et al., 2016); similar representations result in lower decoding accuracies because the classifier confuses them more often than dissimilar representations. As EEG topographies rapidly change over tens to hundreds of milliseconds (Michel and Koenig, 2018), the available information simultaneously changes, leading to changes in decoding accuracy over time. It is not unusual that even before a stimulus is fully presented, the decoding accuracy increases as gradually more information becomes available (see e.g., Crouzet et al., 2015). The resulting time series of decoding accuracies were statistically compared between AP and RP musicians using cluster-based permutation testing (Maris and Oostenveld, 2007).

In detail, for each participant, we resampled the preprocessed and segmented EEG data to 100 Hz using FieldTrip (version 20170713, http://www.fieldtriptoolbox.org/) in MATLAB R2016a. Then, built-in MATLAB functions were used to split and reshape the data to obtain a 32-dimensional vector of amplitudes per trial, per time sample, and per condition. Next, each vector was associated with its corresponding target (listening, labeling). The time sample-wise EEG decoding of the single trials was performed using scikit-learn (version 0.19.2, https://scikit-learn.org/) in Python 3.7.0. Separately for each time sample, we first z-transformed the dataset per feature (i.e. electrode) and subsequently performed the classification of the trials into listening and labeling using a linear Support Vector Machine (C = 1). As some participants had an unequal number of trials in the listening and the labeling condition due to the raw data inspection, we implemented the “balanced” mode of the classifier, in which the weights of the target classes are automatically adjusted to the class frequencies in the data. Decoding accuracy was assessed in a repeated 5-fold stratified cross-validation (100 iterations). Within one iteration, the trials were divided into five folds of approximately equal size, with the restriction that in each fold, the fraction of both target classes was representative of the whole dataset. The classifier was trained on four folds and the remaining fold was used for testing. This procedure was repeated until all folds had been used for testing once.

For the statistical group comparison, the resulting participant-wise time series of decoding accuracies were subsequently analyzed using cluster-based permutation testing in R. First, we calculated a time sample by time sample one-tailed Welch’s t-test to compare decoding accuracies between the groups. Then, a cluster-defining threshold (CDT) of *p* ≤ 0.05 was applied to build clusters of adjacent time samples in which decoding accuracies differed between the groups. Note that this CDT *p* value is used as a descriptive measure and not for inference. The size of the empirical clusters was used as the test statistic. This test statistic was compared to a null distribution of maximum cluster sizes that was obtained by repeating the procedure 10,000 times with permuted group labels. To account for temporal dependencies, we did not permute the group labels of single time samples within a time series of accuracies, but the group labels of every time sample belonging to the same time series were permuted as a whole. This is important because the temporal dependencies within a time series should be preserved, as otherwise the null distribution of cluster sizes is too liberal. The *p* value of the empirical clusters was defined as the fraction of cluster sizes obtained from the permuted data that were larger than the empirical cluster size. The significance level was set to α = 0.05 at cluster level. Please note that the cluster-based permutation test controls the family-wise error rate at the specified α at cluster level.

#### Post-hoc source estimation

To estimate the putative cortical sources underlying differences in time-resolved EEG decoding, we performed source modeling of the EEG signal using Brainstorm (version 3.4; https://neuroimage.usc.edu/brainstorm/) (Tadel et al., 2011). We used default Brainstorm settings unless otherwise stated. First, we computed a head model based on the default ICBM152 anatomical template with 15,002 dipole locations using the OpenMEEG boundary element method (Gramfort et al., 2010) and calculated the noise covariance matrix based on the baseline (−100–0 ms) of the single trials. Then, we estimated distributed sources for each participant and condition using minimum norm estimation with loosely constrained dipole orientations (1 normal to cortex dipole, 2 tangential dipoles with amplitude = 0.2).

### Time-frequency-resolved single-trial EEG decoding

The time-frequency-resolved EEG decoding was in large parts identical to the time-resolved EEG decoding. The crucial difference was the transformation of the time-domain signal to the frequency domain, which enabled frequency-band-specific EEG decoding to study the neural representations underlying tone listening and labeling. In detail, we first applied time-frequency analysis to the preprocessed EEG data to calculate the frequency-specific power over time (Cohen, 2014). The time-frequency analysis was performed using the FieldTrip method *mtmconvol* in combination with a single Hanning taper. The power values were calculated in a sliding time window of 500 ms (fixed length over frequencies) which was moved in 50 ms steps. For fixed window lengths, the frequency resolution (in Hz) is determined by the inverse of the window length (in s); this resulted in a frequency resolution of 2 Hz. We excluded frequencies below 4 Hz and above 30 Hz from the analysis because for frequencies in the delta range (below 4Hz), a 500 ms time window includes only one cycle, and for frequencies in the gamma range (above 30 Hz), there is a high possibility for contamination by muscle artifacts (Whitham et al., 2007). Consequently, we analyzed the following frequency bands: theta = 4–6 Hz, alpha = 8–12 Hz, and beta = 14–30 Hz. We then used built-in MATLAB functions to split and reshape the data to obtain a vector of power values per trial, per time sample, per condition, and per frequency band (theta, alpha, beta). We again associated each vector with its target (listening, labeling) and, separately for each frequency band, performed the time sample-wise single-trial EEG decoding using scikit-learn. This resulted in participant-wise time series of decoding accuracies, which were subjected to cluster-based permutation testing in R, separately for each frequency band. See above for details concerning the EEG decoding and permutation testing.

### Univariate analyses of the EEG data

To keep the study comparable with previous EEG studies on AP and RP, we additionally performed univariate statistics on the EEG data in both the time domain and the frequency domain. For the univariate analysis in the time domain, we resampled the data to 100 Hz. We computed ERPs by averaging the single trials per participant and condition. To analyze the interaction between group and condition, we calculated difference waves (labeling minus listening) for each participant. Based on the grand average of these difference waves across both groups, we restricted the analysis to the time interval between 300 and 600 ms relative to stimulus onset (cf. Figure 4A). We compared the difference waves between the groups using cluster-based permutation testing across both time and electrodes in FieldTrip (10,000 permutations, independent samples t-test, CDT *p* ≤ 0.05, α = 0.05 at cluster level; minimum number of electrodes for cluster = 4); in case of a significant interaction, we performed follow-up analyses by repeating the group comparison within each condition separately, and appropriately adjusted the significance level to α = 0.025. To compare the univariate analysis in the time domain with the time-resolved EEG decoding, we repeated the group comparisons without restricting the statistical analysis to a predefined time interval (analysis window = 0– 800 ms).

An analogous procedure was employed for the univariate analysis in the frequency domain. We averaged the single-trial time-frequency power to compute an average time-frequency representation per participant and condition. To analyze the group x condition interaction, we calculated the participant-wise difference between the conditions (labeling minus listening) in time-frequency power. Based on the grand average of the differences in time-frequency power across all participants, we restricted the analysis to the theta frequency band and the time interval between 300 and 800 ms (cf. Figure 5A). Finally, we performed cluster-based permutation testing on the differences in time-frequency power across time, frequency, and electrodes in FieldTrip (10,000 permutations, independent samples t-test, CDT *p* ≤ 0.05, α = 0.05 at cluster level); in case of a significant interaction, we performed follow-up group comparisons within each condition separately (α = 0.025). Again, for a comparison with the time-frequency-resolved EEG decoding, we repeated the group comparisons for each of the three frequency bands (theta, alpha, and beta) without restricting the statistical analysis to a predefined time interval.

## Results

### Behavioral results

Descriptive statistics of the participants’ demographical and behavioral characteristics are given in Table 1. Group comparison of these characteristics revealed that there was no statistically significant difference in age (*t*(100.97) = 1.33, *p* = 0.19, *d* = 0.26), age of onset of musical training (*t*(102.42) = −1.20, *p* = 0.23, *d* = 0.23), cumulative musical training (*t*(99.71) = 1.43, *p* = 0.16, *d* = 0.28), crystallized intelligence (*t*(102.86) = −1.49, *p* = 0.14, *d* = 0.29), and fluid intelligence (*t*(100.82) = −1.54, *p* = 0.13, *d* = 0.30). We found a significant but nevertheless small difference in musical aptitude as measured by the AMMA total score (*t*(100.99) = 2.14, *p* = 0.03, *d* = 0.42), which was driven by higher AMMA tonal scores in AP musicians (*t*(100.61) = 2.44, *p* = 0.02, *d* = 0.48); the AMMA rhythm scores did not significantly differ (*t*(101.38) = 1.53, *p* = 0.13, *d* = 0.30). As shown in the left panel of Figure 1, we found a substantially better tone-naming proficiency in AP musicians (*t*(102.92) = 13.94, *p* < 2.2 × 10^−16^, *d* = 2.72). In the Stroop-like task, we found a significantly larger Stroop-effect in AP musicians associated with a medium effect size (*t*(102.65) = 2.78, *p* = 0.007, *d* = 0.54), which confirmed the presence of labeling automaticity in our sample of AP musicians (see Figure 1, right panel). There was no significant group difference in the interval identification score (*t*(86.53) = 1.18, *p* = 0.24, *d* = 0.23).

**Table 1.** Participant characteristics. Continuous measures given as mean ± standard deviation.

|  | AP musicians | RP musicians |
| --- | --- | --- |
| Number of participants | 54 | 51 |
| Sex (female / male) | 27 / 27 | 24 / 27 |
| Handedness (right / left / both) | 47 / 4 / 3 | 46 / 4 / 1 |
| Age | 26.67 ± 5.49 years | 25.37 ± 4.49 years |
| Musical aptitude (AMMA) – total | 66.11 ± 6.31 | 63.35 ± 6.86 |
| Musical aptitude (AMMA) – tonal | 32.33 ± 3.75 | 30.45 ± 4.13 |
| Musical aptitude (AMMA) – rhythm | 33.78 ± 2.83 | 32.90 ± 3.03 |
| Age of onset of musical training | 5.93 ± 2.39 | 6.49 ± 2.44 |
| Cumulative musical training | 16,592.51 ± 12,245.67 hours | 13,539.00 ± 9,609.84 hours |
| Crystallized intelligence (MWT-B) | 27.69 ± 5.10 | 29.10 ± 4.64 |
| Fluid intelligence (KAI) | 123.41 ± 32.16 | 132.19 ± 26.16 |
| Tone-naming score | 76.41 ± 19.55 % | 24.00 ± 18.97 % |
| Stroop-effect | 39.32 ± 38.71 ms | 18.32 ± 38.74 ms |
| Interval identification score | 77.59 ± 15.94 % | 73.00 ± 22.97 % |

Abbreviations: AMMA = Advanced Measures of Music Audiation, AP = absolute pitch, KAI = Kurztest für allgemeine Basisgrößen der Informationsverarbeitung, MWT-B = Mehrfachwahl-Wortschatz-Intelligenztest, RP = relative pitch.

**Figure 1.**
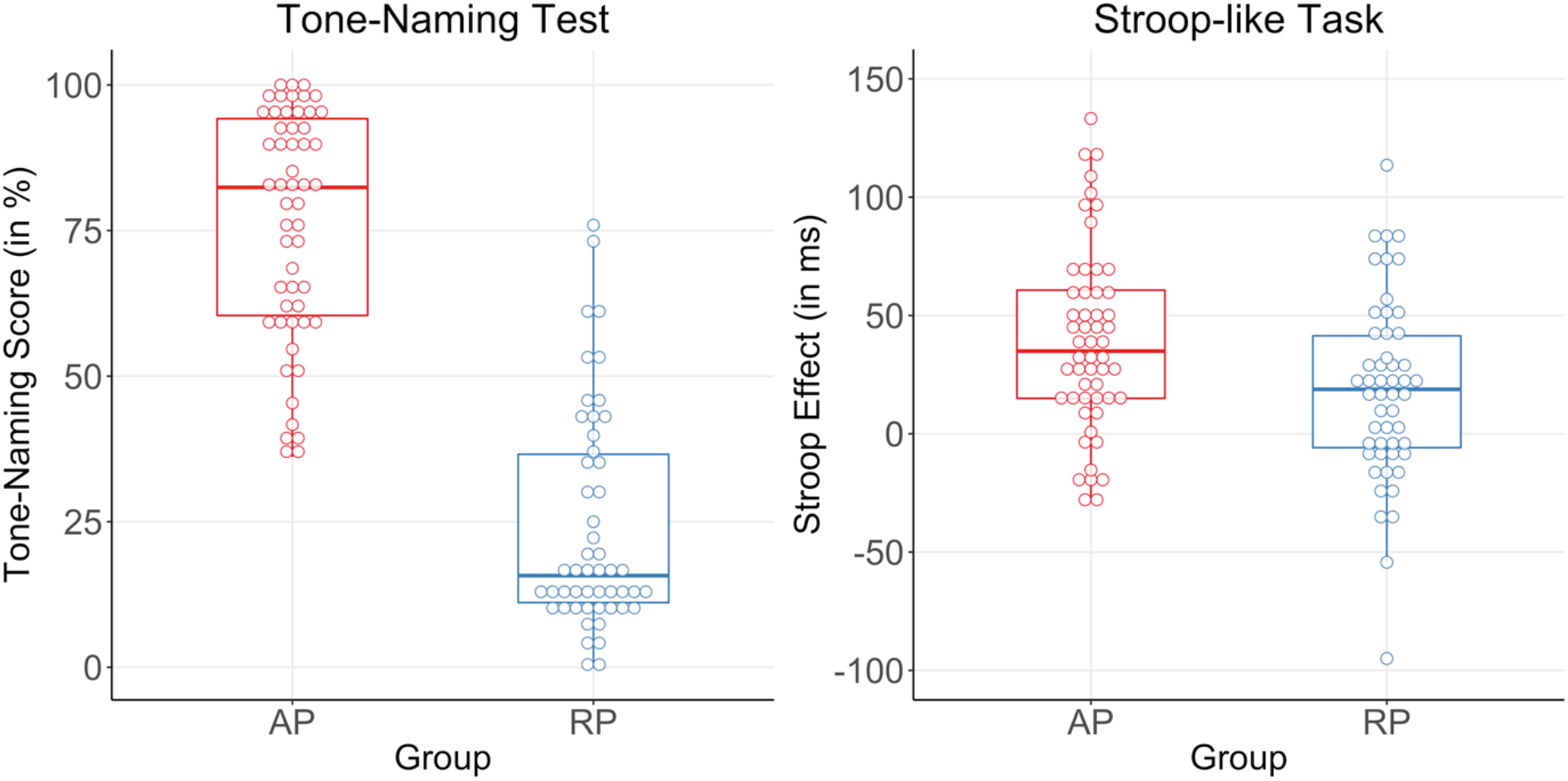
Results of the tone-naming test and the Stroop-like task. The tone-naming test revealed a substantially better tone-naming proficiency in AP musicians than RP musicians (left panel). The Stroop-like task confirmed the presence of labeling automaticity in our sample as AP musicians showed a larger Stroop-effect than RP musicians (right panel). Abbreviations: AP = absolute pitch, RP = relative pitch.

### Exploratory correlation analyses among behavioral characteristics

As an association between tone naming and interval identification has been reported previously (Dooley and Deutsch, 2011), we tested the presence of this association in our sample using Pearson correlations. Across the whole sample, tone-naming proficiency and interval-identification scores were significantly correlated (*r* = 0.31, *p* = 0.002), and this association was also present within the groups of AP musicians (*r* = 0.32, *p* = 0.02) and RP musicians (*r* = 0.40, *p* = 0.004). These correlations were of medium size according to Cohen (1992). We also correlated the tone-naming proficiency with the size of the Stroop-effect to check whether the precision of tone naming was associated with the automaticity of labeling. We found a correlation between tone-naming and Stroop-effect across the whole sample (*r* = 0.24, *p* = 0.02), but this correlation was likely driven by group differences in both measures as we found no significant correlations within the groups (AP: *r* = −0.12, *p* = 0.37; RP: *r* = 0.22, *p* = 0.13).

### Results of the time-resolved EEG decoding

The participant-wise time-resolved EEG decoding of listening and labeling revealed that in the early stages of pitch perception, decoding accuracies were near chance level (i.e. the representational similarity of the conditions was high) in many participants of both groups. In later stages, decoding accuracies increased before becoming lower again during very late stages of pitch perception. See Supplementary Figure 1A for a participant-wise visualization of time-resolved EEG decoding and Supplementary Figure 1B for a participant-wise visualization of statistical significance of time-resolved EEG decoding. The group comparison of the decoding time series using cluster-based permutation testing yielded a significant cluster in the time interval between 430 and 550 ms after stimulus onset (*p* = 0.01, empirical cluster size *k* = 13). Consistent with our predictions, EEG decoding was significantly lower in AP musicians and thus, the neural representations underlying tone listening and labeling were more similar. During and immediately after stimulus presentation (~0–400 ms), we found no evidence for a difference in decoding accuracies between the groups. Figure 2A visualizes the group-wise decoding time series, and Figure 2B visualizes the null distribution of cluster sizes (obtained through 10,000 permutations) of the time-resolved analysis.

**Figure 2.**
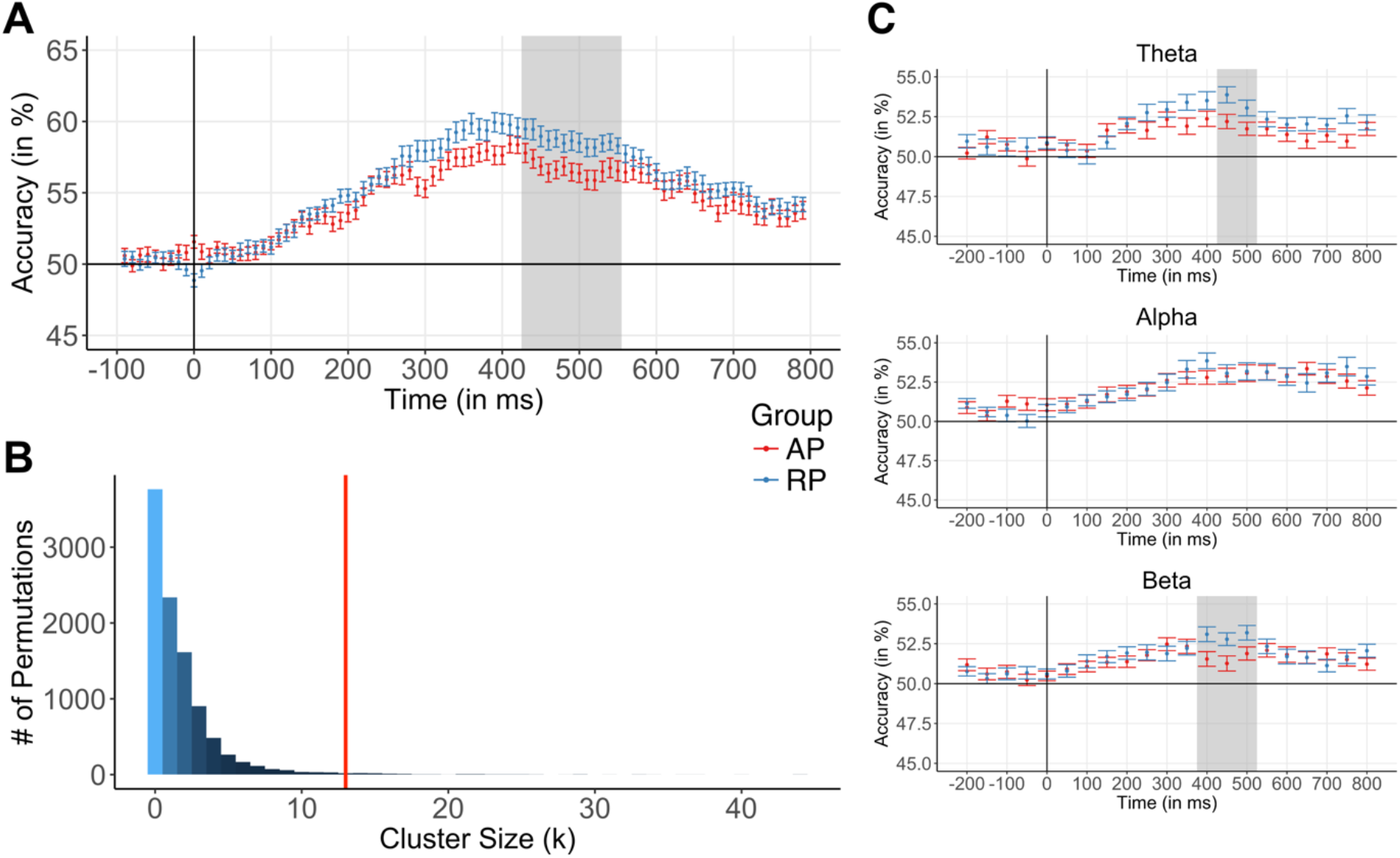
Single-trial EEG decoding in the time and frequency domain. Group differences in the representational similarity of listening and labeling were found both in the time domain and the frequency domain. (A) Time-resolved EEG decoding revealed group differences in EEG decoding 430–550 ms after stimulus onset (gray shaded area). (B) The null distribution of cluster sizes was obtained by repeating the analysis 10,000 times with permuted group labels. The empirical cluster size (k = 13) is depicted by the red vertical line. (C) Time-frequency-resolved EEG decoding revealed group differences in representational similarity in the theta frequency band 450–500 ms after stimulus onset (gray shaded area in the upper panel) and the beta frequency band 400–500 ms after stimulus onset (gray shaded area in the lower panel). In the alpha frequency band, EEG decoding did not differ between the groups. Error bars represent the standard error of the mean per time sample and group. Abbreviations: AP = absolute pitch, RP = relative pitch.

#### Results of the post-hoc source estimation

We performed source estimation for the time interval where we found significant group differences in representational similarity (430–550 ms). For each condition separately, we averaged the source-space current density values across time between 430 and 550 ms using the absolute values of the three dipole orientations (1 normal, 2 tangential). Subsequently, we subtracted the current density values in the listening condition from the values in the labeling condition and averaged these differences per group. As shown in Figure 3, in both groups, maximal condition differences in source space were located in the presupplementary motor area (preSMA), and, to a lesser degree, in the medial superior parietal cortex and the dorsolateral prefrontal cortex. Descriptively, condition differences in source space were stronger in RP musicians than AP musicians. We refrained from additional statistical inference in source space as this would constitute a circular analysis, given that we identified the time window based on significant group differences in time-resolved EEG decoding.

**Figure 3.**
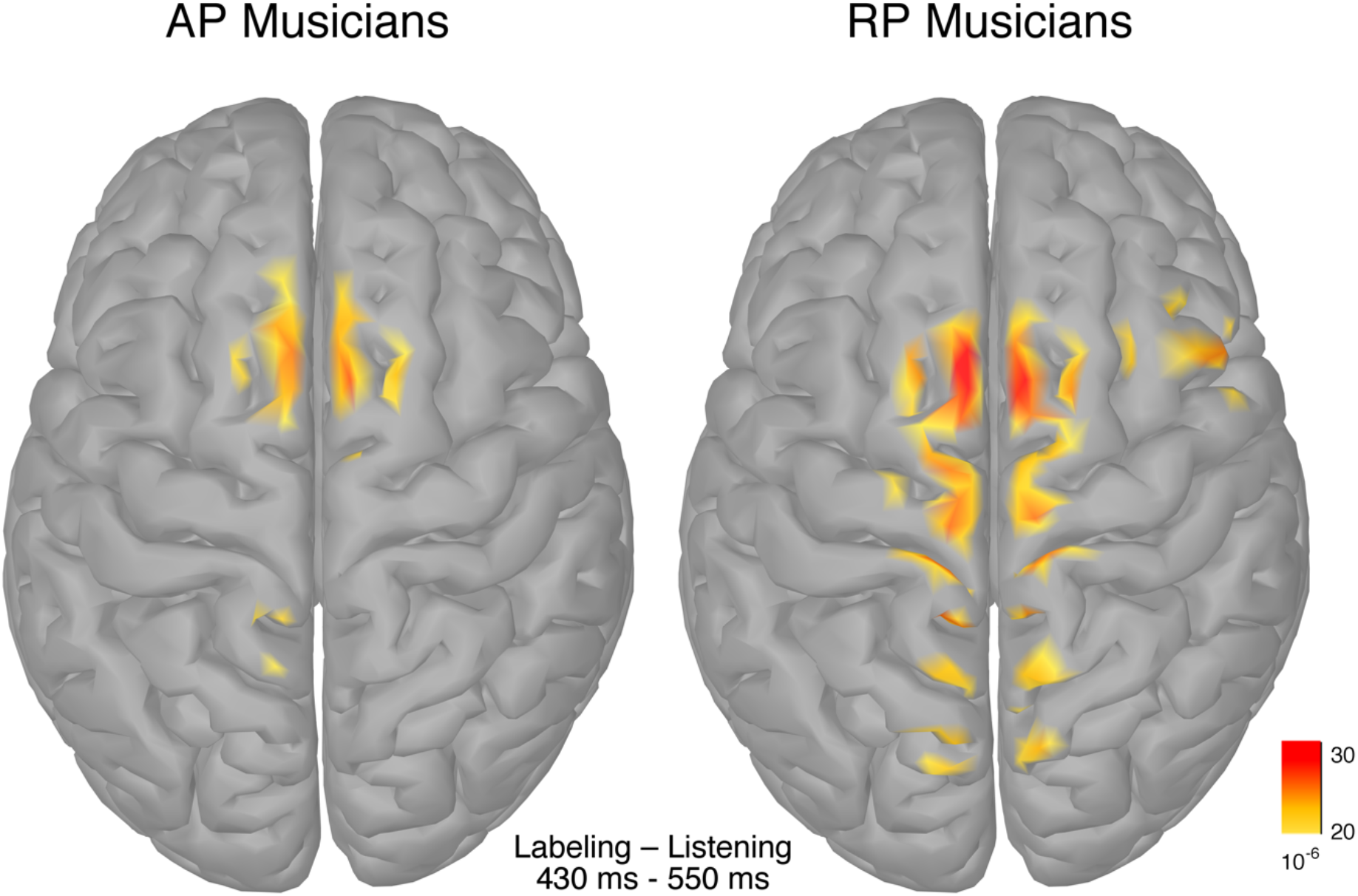
Group-specific condition differences in source space. The group-specific condition differences (labeling minus listening) for the time interval where representational similarity of listening and labeling differed between the groups (430–550 ms) were located in the presupplementary motor area, and, to a lesser degree, in the medial superior parietal cortex and the dorsolateral prefrontal cortex. For visualization purposes, the images were thresholded at 60% of the maximal current density amplitude.

### Results of the time-frequency-resolved EEG decoding

The participant-wise time-frequency-resolved EEG decoding of the experimental conditions revealed that in the theta and beta frequency bands, for many participants, decoding accuracies were relatively low during the first stages of pitch perception (i.e. the neural representations were similar), gradually increased during later stages and then dropped again in the very late stages (cf. Supplementary Figures 2A, 2B, 3A, and 3B). The group comparison of the time-frequency-resolved decoding time series using cluster-based permutation testing revealed significant group differences in EEG decoding in the theta and the beta frequency bands. Analogous to the time-resolved analysis, in the theta frequency band, we found significantly lower decoding accuracies in AP musicians between 450 and 500 ms after stimulus onset (*p* = 0.04, *k* = 2). In the beta frequency band, we found lower decoding accuracies in AP musicians between 400 and 500 ms (*p* = 0.02, *k* = 3). In contrast, we did not find evidence for differences in EEG decoding in the alpha frequency band. Figure 2C visualizes the group-wise decoding time series for each frequency band.

### Results of the univariate analyses

Using a restricted analysis window, the univariate analysis of the time-domain EEG data revealed a group (AP vs. RP) x condition (listening vs. labeling) interaction 380–530 ms after stimulus onset (*p* = 0.02, *k* = 16), characterized by smaller difference wave amplitudes (labeling minus listening) in AP musicians at predominantly frontal, central, and parietal electrodes (see Figure 4B and Figure 4C). Post-hoc group comparisons separately within each condition revealed no significant amplitude differences (all *p* > 0.025). However, there was one cluster of lower amplitudes in AP musicians during labeling, which did not survive the cluster-based correction for multiple comparisons (*p* = 0.11, *k* = 3). From this it follows that in the time domain, the groups could primarily be differentiated based on their condition differences; the group difference within each condition, though, was too small to be reliably detected. The univariate time-domain analysis without a restriction of the temporal analysis window revealed the same interaction as the restricted analysis (*p* = 0.03, *k* = 16).

**Figure 4.**
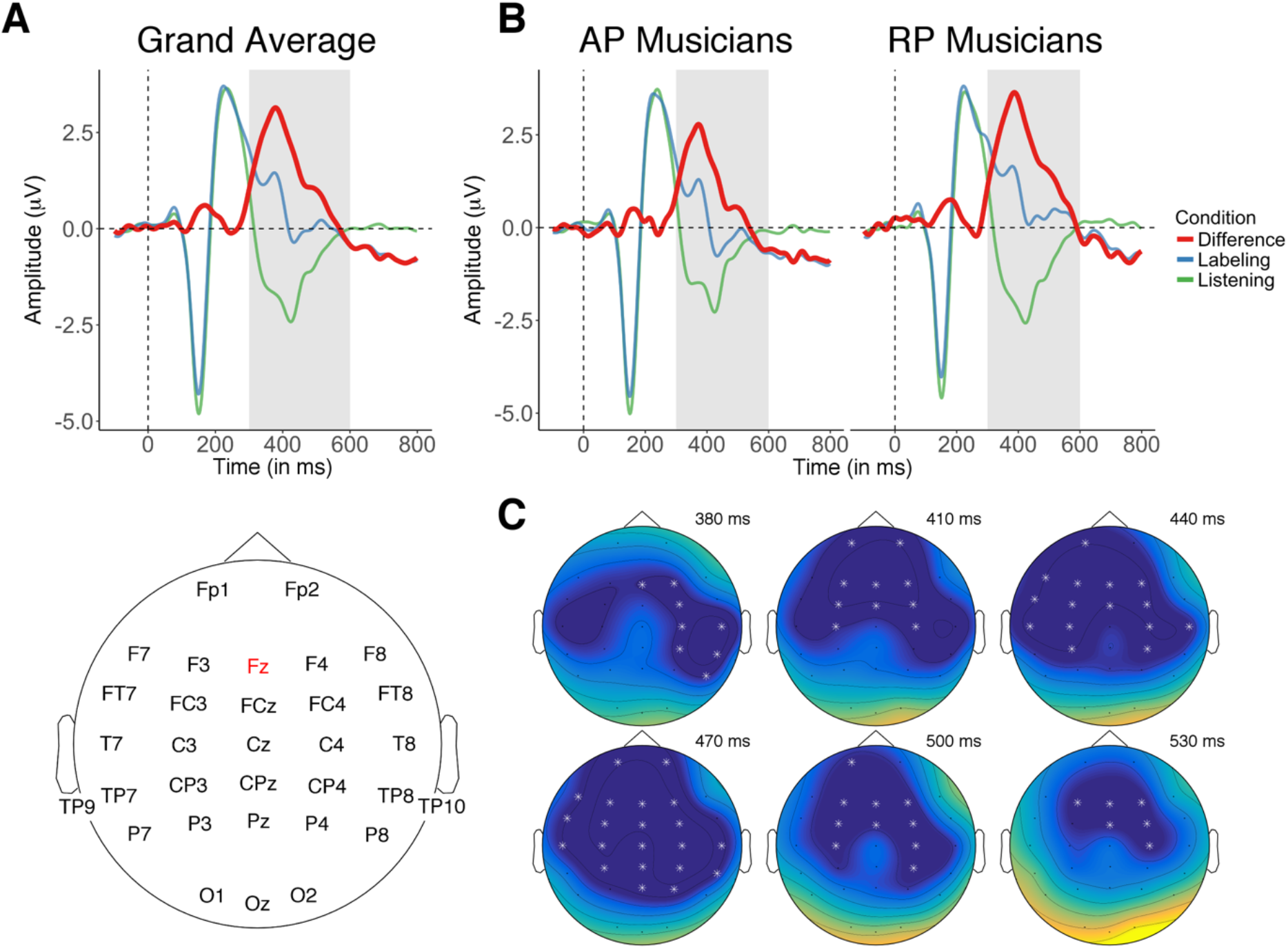
Univariate analysis of the time-domain EEG data. (A) Based on the grand-average difference wave of all participants (in red) at electrode Fz, we restricted the cluster-based permutation testing to 300–600 ms after stimulus onset (gray shaded area). We also analyzed the time-domain EEG data without this restriction (see main text). (B) Waveforms per condition (in blue and green) and difference waveform (in red), separately for AP musicians (left) and RP musicians (right). (C) Using a restricted analysis window, cluster-based permutation testing revealed reduced difference wave amplitudes (labeling minus listening) in AP musicians at predominantly frontal, central, and parietal electrodes. The highlighted electrodes showed the strongest group differences.

Using a restricted analysis window, the univariate analysis in the frequency domain revealed a significant group x condition interaction in the theta frequency band 450–750 ms after stimulus onset (*p* = 0.04, *k* = 7); this interaction was characterized by smaller power differences between conditions (labeling minus listening) in AP musicians (see Figure 5B). The post-hoc analyses separately within each condition revealed no group differences during listening but smaller theta power in AP musicians during labeling (*p* = 0.003, *k* = 11). The univariate frequency-domain analysis without restriction of the analysis window only yielded a trend towards a group x condition interaction (*p* = 0.05, *k* = 7). Furthermore, the univariate frequency-domain analysis did not reveal significant group differences in the alpha and beta frequency bands without a restriction of the analysis window.

**Figure 5.**
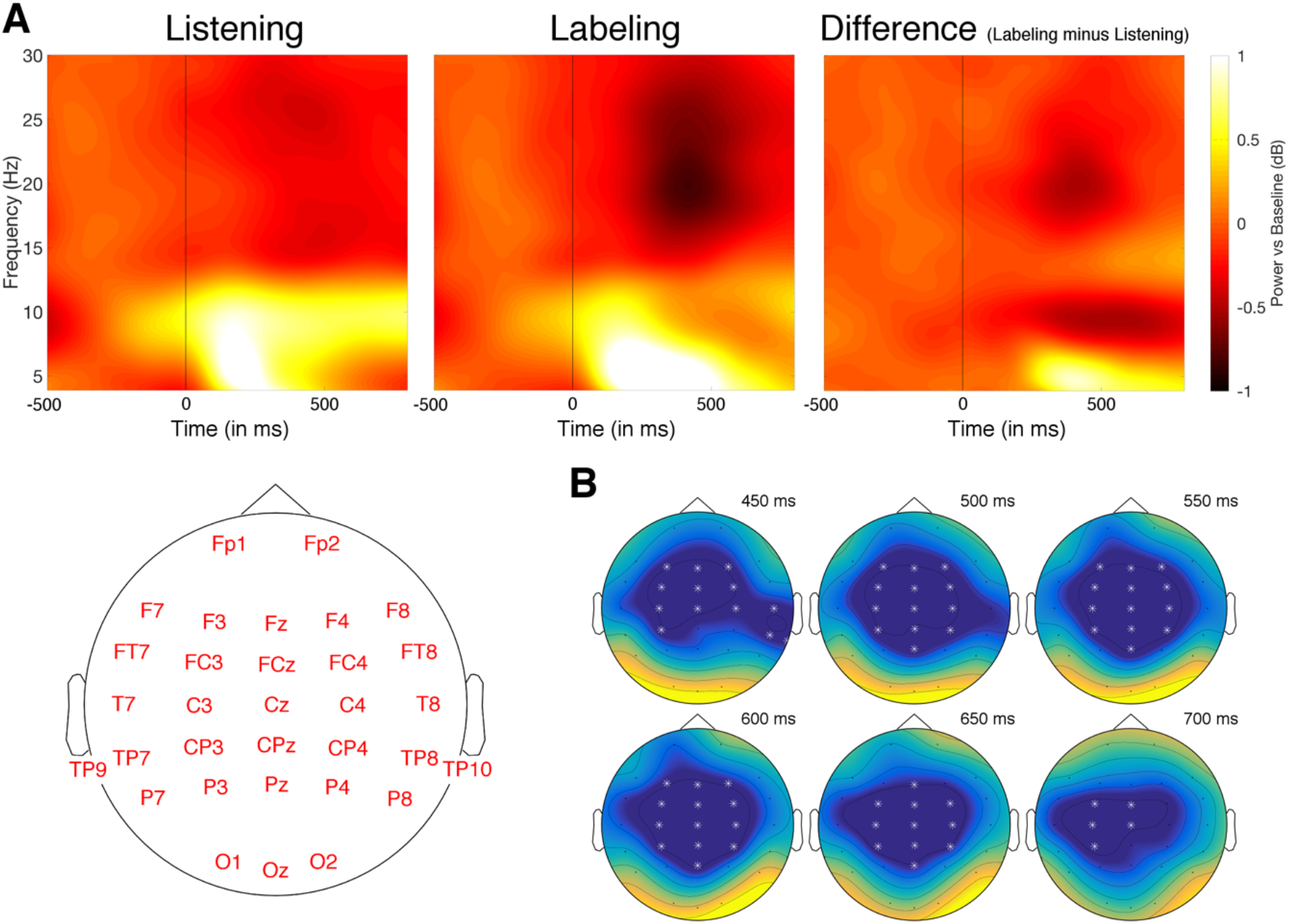
Univariate analysis of the frequency-domain EEG data. (A) Based on the grand average of differences in time-frequency power across all participants (right panel), we restricted the cluster-based permutation testing to the theta frequency band (4–6 Hz) and the time interval between 300 and 800 ms after stimulus onset. We also analyzed the frequency-domain EEG data without this restriction (see main text). The mean time-frequency power over all electrodes is visualized. (B) The restricted cluster-based permutation testing revealed reduced theta power differences (labeling minus listening) in AP musicians. The highlighted electrodes showed the strongest group differences in the theta frequency band.

## Discussion

The conceptualization of multivariate EEG patterns as decodable neural representations is a powerful tool for investigating the dynamic neural mechanisms underlying perception and cognition (Grootswagers et al., 2016). As explained in the introduction, both cognitive models of AP and empirical evidence from previous studies suggest more similar representations of tone listening and labeling in AP musicians than in RP musicians. However, no study has explicitly tested this prediction. Here, we studied the neural representations underlying listening and labeling in a large sample of AP and RP musicians using a “brain-decoding” approach. As predicted, time-resolved single-trial EEG decoding revealed higher representational similarity of listening and labeling in AP musicians during late stages of pitch perception. Time-frequency-resolved single-trial EEG decoding revealed that the higher representational similarity in AP musicians was present in oscillations in the theta and beta frequency bands. In the alpha frequency band, we found no evidence for group differences with regard to the similarity of neural representations underlying listening and labeling. Univariate analysis in the time domain revealed that AP musicians had lower amplitude differences between the conditions. Frequency-domain univariate analysis revealed lower theta power differences in AP musicians driven by lower theta during labeling. In contrast to the decoding approach, the univariate approach did not identify group differences in the beta frequency band.

The dynamics of representational similarity were characterized by a lack of differences between AP and RP musicians during the early stages of pitch perception; only after 300 ms, group differences began to emerge, and these differences were strongest during late processing stages. The early processing of tones encompasses the perceptual analysis of acoustic features of the stimulus (Koelsch and Siebel, 2005); hence, our results suggest that AP and RP musicians do not differ in this initial analysis of the tones. This is in accordance with a number of previous EEG studies not finding differences between AP and RP musicians in early ERPs (Elmer et al., 2013; Greber et al., 2018; Pantev et al., 1998; Rogenmoser et al., 2015; Tervaniemi et al., 1993). On the other hand, much evidence points to differences between AP and RP musicians in late ERPs associated with higher-order cognitive functions (Bischoff Renninger et al., 2003; Crummer et al., 1994; Elmer et al., 2013; Hantz et al., 1992; Klein et al., 1984; Wayman et al., 1992). We extend these studies by demonstrating that by using a decoding approach, group differences are detectable on a single-trial level. Consistent with the notion that late stages of pitch perception are associated with cognitive processing, the putative cortical sources underlying the group differences were located in the superior frontal and parietal cortices.

In previous EEG and neuroimaging studies, AP and RP have been repeatedly associated with tonal working memory, e.g., in the context of auditory oddball paradigms (Hantz et al., 1992; Klein et al., 1984). The original idea was that RP musicians need to update working memory when being confronted with novel incoming tones, whereas AP musicians do not rely on working memory processes during pitch perception because they have fixed long-term memory templates for tones (Klein et al., 1984). This idea has been further developed to suggest that AP musicians do not need to access working memory to name musical intervals (Hantz et al., 1992; Zatorre et al., 1998) or to label single tones (Itoh et al., 2005); the automaticity of tone labeling as demonstrated by our Stroop-like task underlines this observation. Furthermore, in recent behavioral studies, tone-naming proficiency has been linked to working memory capacity (Deutsch and Dooley, 2013; Van Hedger et al., 2015). Taken together, in the case of our pitch perception task, the dissimilarity of the neural representations underlying listening and labeling in RP musicians might be due to them accessing working memory during labeling (but not during listening).

To date, only a single study has investigated the neural oscillations of AP and RP during musical perception (Behroozmand et al., 2015). Using univariate analysis, the authors did not identify differences between AP and RP musicians. In this study, we found group differences in theta and beta oscillations in late stages of pitch perception. The exact role of neural oscillations in perception and cognition is still somewhat of a mystery (Wang, 2010), but both theta and beta oscillations have been associated with various cognitive functions. In the auditory domain, it has been proposed that theta oscillations have a causal role in enhancing tonal working memory (Albouy et al., 2017). Beta oscillations have been linked to top-down processing (Engel and Fries, 2010).

However, as we did not collect behavioral measurements of higher-order cognitive functions apart from the Stroop-like task, the specific nature of the cognitive functions underlying the group differences in late stages of pitch perception remains speculative. Future studies should include such behavioral measurements to better determine the underlying cognitive processing.

As a supplement to the EEG decoding approach, we used classical univariate analysis, first and foremost to make our study comparable with previous ERP studies on AP. We found, on average, lower amplitude differences and, using a restricted analysis window, lower theta power differences in AP musicians. These results are somewhat comparable to the results of the EEG decoding analysis. However, using the same analysis windows for both approaches, the decoding approach showed higher statistical sensitivity in the frequency domain as it exclusively identified a group difference in the beta frequency band. In addition, the univariate analysis without a restricted analysis window only revealed a trend towards lower theta power differences in AP musicians. This is consistent with observations regarding a higher sensitivity of multivariate compared to univariate methods in detecting subtle distributed effects (Grootswagers et al., 2016). Therefore, one should not mistakenly equate the results of the two analyses approaches; they presumably reflect two complementary aspects of the same underlying neural mechanisms, but the relationship of these two aspects is still subject to ongoing discussions (Jimura and Poldrack, 2012). In the case of EEG, a major advantage of (time-resolved) decoding is that it summarizes the information from multiple experimental conditions, electrodes, and trials into a single time series that can potentially be related to signals from other imaging modalities, e.g., neuroimaging data, behavioral data, or even the outputs of computational models (Kriegeskorte et al., 2008). In this context, future studies investigating the neural representations underlying pitch perception should try to sample more than two conditions (listening, labeling) in the vast space of possible experimental conditions to fully exploit the capabilities of this approach. Finally, it should be noted that apart from representational similarity analysis as it was performed here, there exist other analysis methods that allow for a multivariate analysis of EEG patterns, e.g., the multivariate general linear model (Friston et al., 1996).

In conclusion, we showed that the neural representations underlying listening and labeling are more similar in AP musicians than RP musicians during late stages of pitch perception; this effect was present in oscillations in the theta and beta frequency bands. Compared to the novel decoding approach, the conventional univariate approach was less sensitive in identifying subtle group differences in the frequency domain.

## Acknowledgements

This work was supported by the Swiss National Science Foundation (SNSF), grant no. 320030_163149 to LJ. We thank our research interns Anna Speckert, Chantal Oderbolz, Fabian Demuth, Florence Bernays, Joëlle Albrecht, Kathrin Baur, Laura Keller, Melek Haçan, Nicole Hedinger, Pascal Misala, Petra Meier, Sarah Appenzeller, Tenzin Dotschung, Valerie Hungerbühler, Vanessa Vallesi, and Vivienne Kunz for their invaluable help in data acquisition and research administration. Furthermore, we thank Christian Brauchli and Anja Burkhard for their support within the larger absolute pitch project, and Carina Klein, Stefan Elmer, and all the members of the Auditory Research Group Zurich (ARGZ) for their valuable comments on the experimental procedure.

## Conflicts of Interest

The authors declare no conflicts of interest.

## Supplemental Files

**Supplementary Figure 1A.**
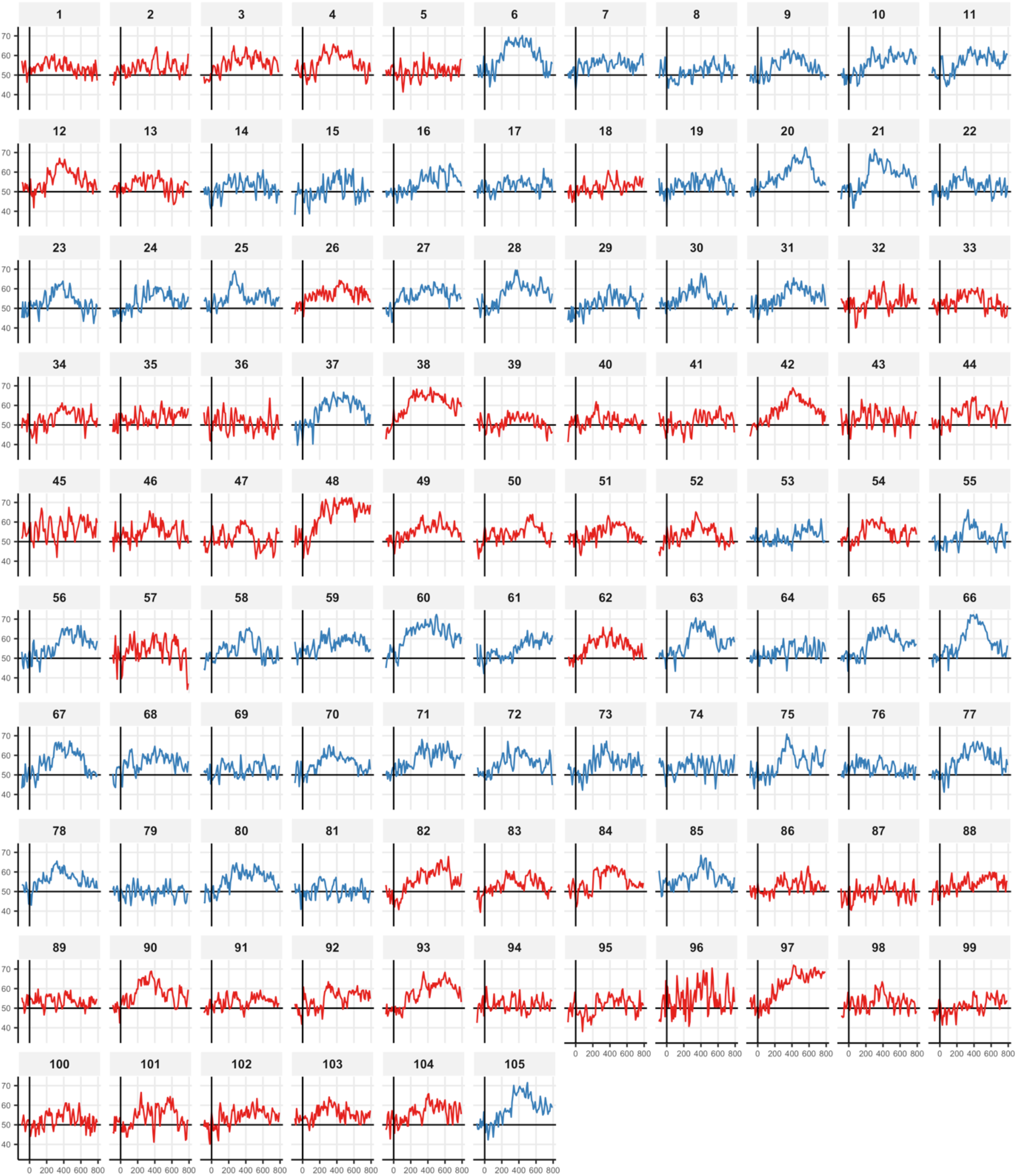
Participant-wise decoding time series of the time-resolved EEG decoding. The x-axis represents time (in ms) relative to stimulus onset; the y-axis represents decoding accuracy (in %). The solid black vertical line denotes stimulus onset. The solid black horizontal line at y = 50 denotes chance level (50%). Absolute pitch musicians are colored in red; relative pitch musicians are colored in blue.

**Supplementary Figure 1B.**
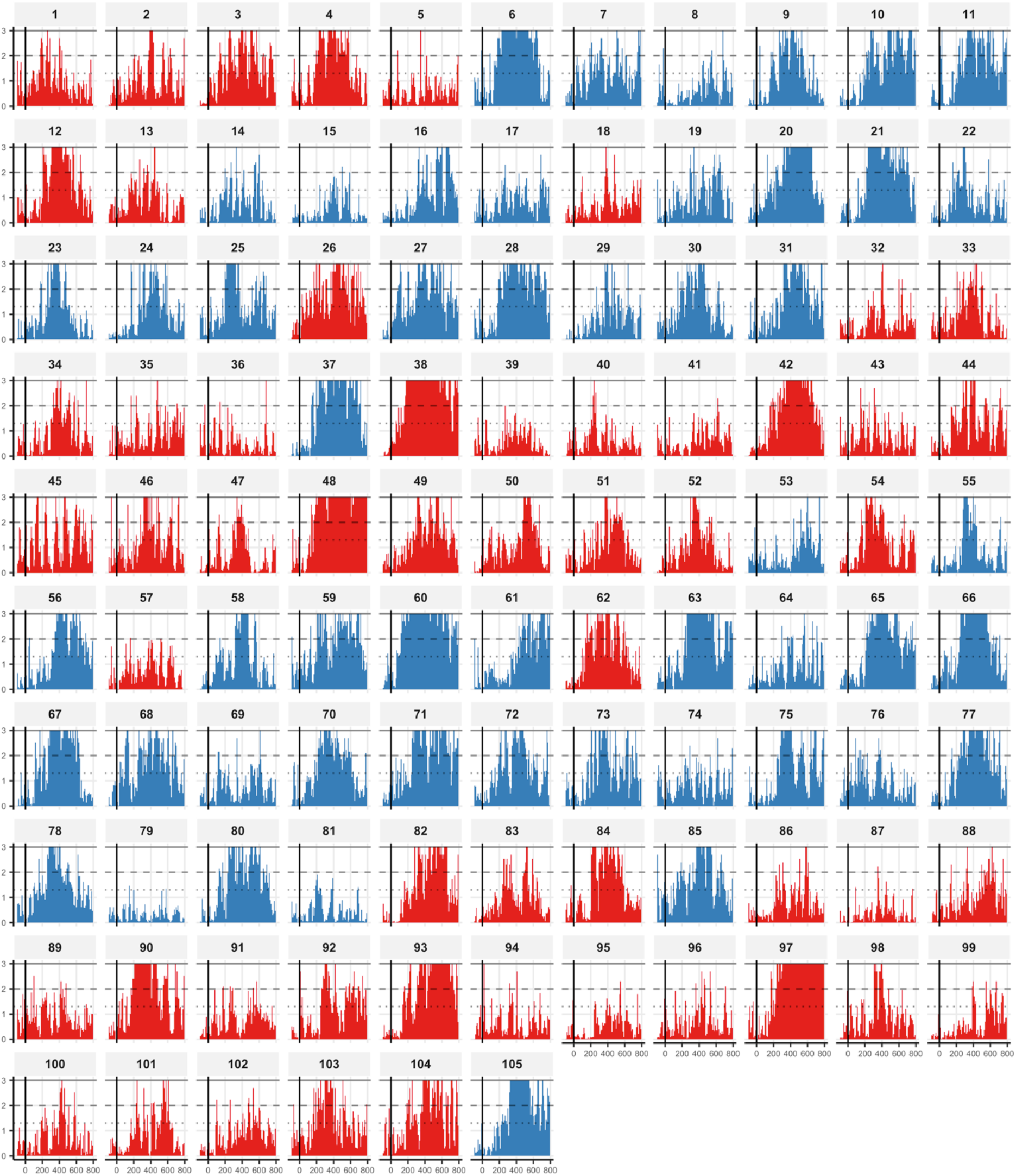
Participant-wise −log_10_ *p* value time series of the time-resolved EEG decoding. Group differences in representational similarity (operationalized by decoding accuracy) are unlikely to be meaningful if the accuracies in both groups are below chance level (50%). Thus, we assessed the statistical significance of EEG decoding for each participant and time bin separately using 1,000 random permutations. The x-axis represents time (in ms) relative to stimulus onset; the y-axis represents −log_10_ *p* values. The solid black vertical line denotes stimulus onset. The dotted horizontal line at y ≈ 1.3 denotes *p* = 0.05, the dashed horizontal line at y = 2 denotes *p* = 0.01, and the solid horizontal line at y = 3 denotes *p* = 0.001. Absolute pitch musicians are colored in red; relative pitch musicians are colored in blue.

**Supplementary Figure 2A.**
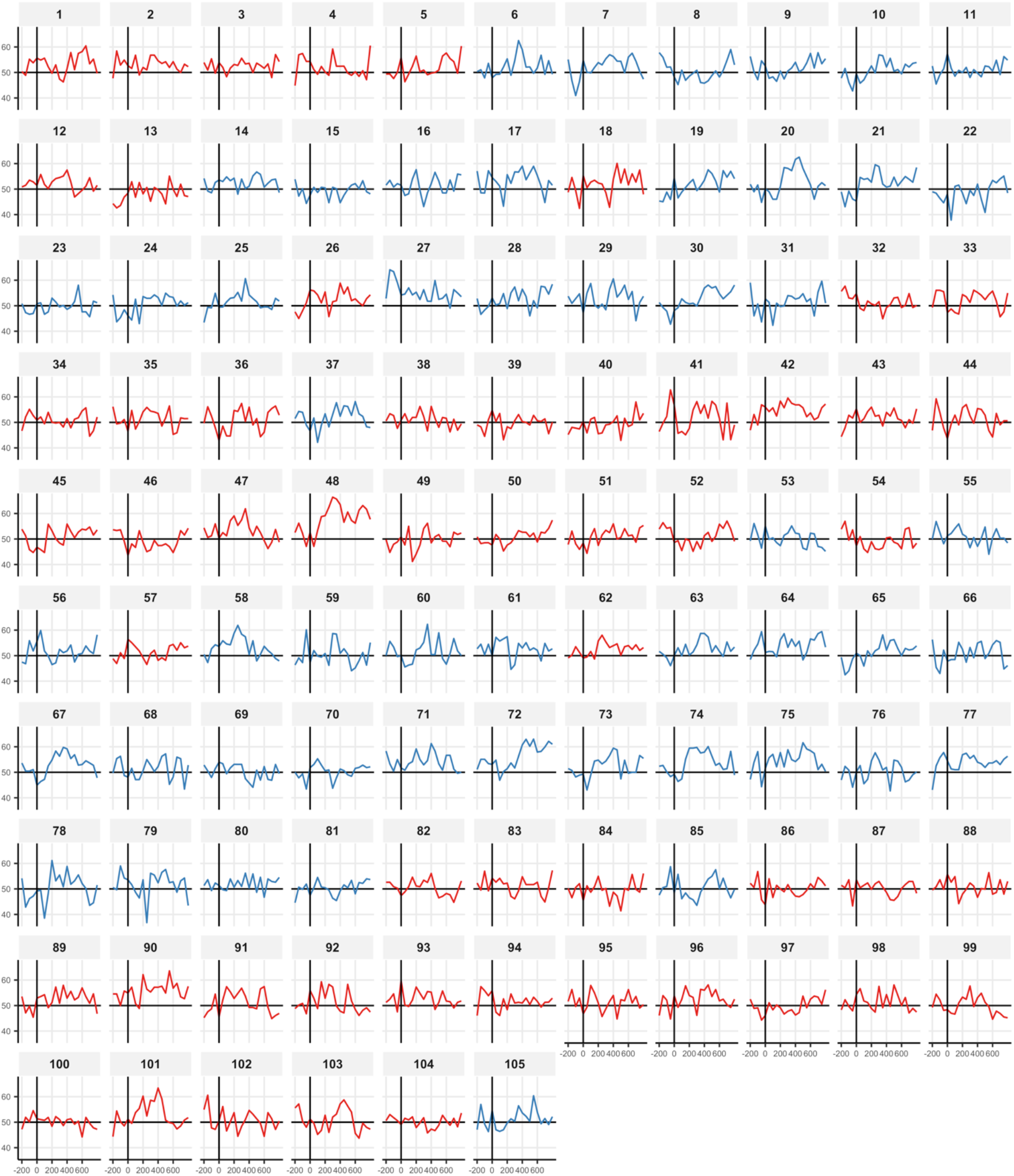
Participant-wise decoding time series of the time-frequency-resolved EEG decoding in the theta frequency band. The x-axis represents time (in ms) relative to stimulus onset; the y-axis represents decoding accuracy (in %). The solid black vertical line denotes stimulus onset. The solid black horizontal line at y = 50 denotes chance level (50%). Absolute pitch musicians are colored in red; relative pitch musicians are colored in blue.

**Supplementary Figure 2B.**
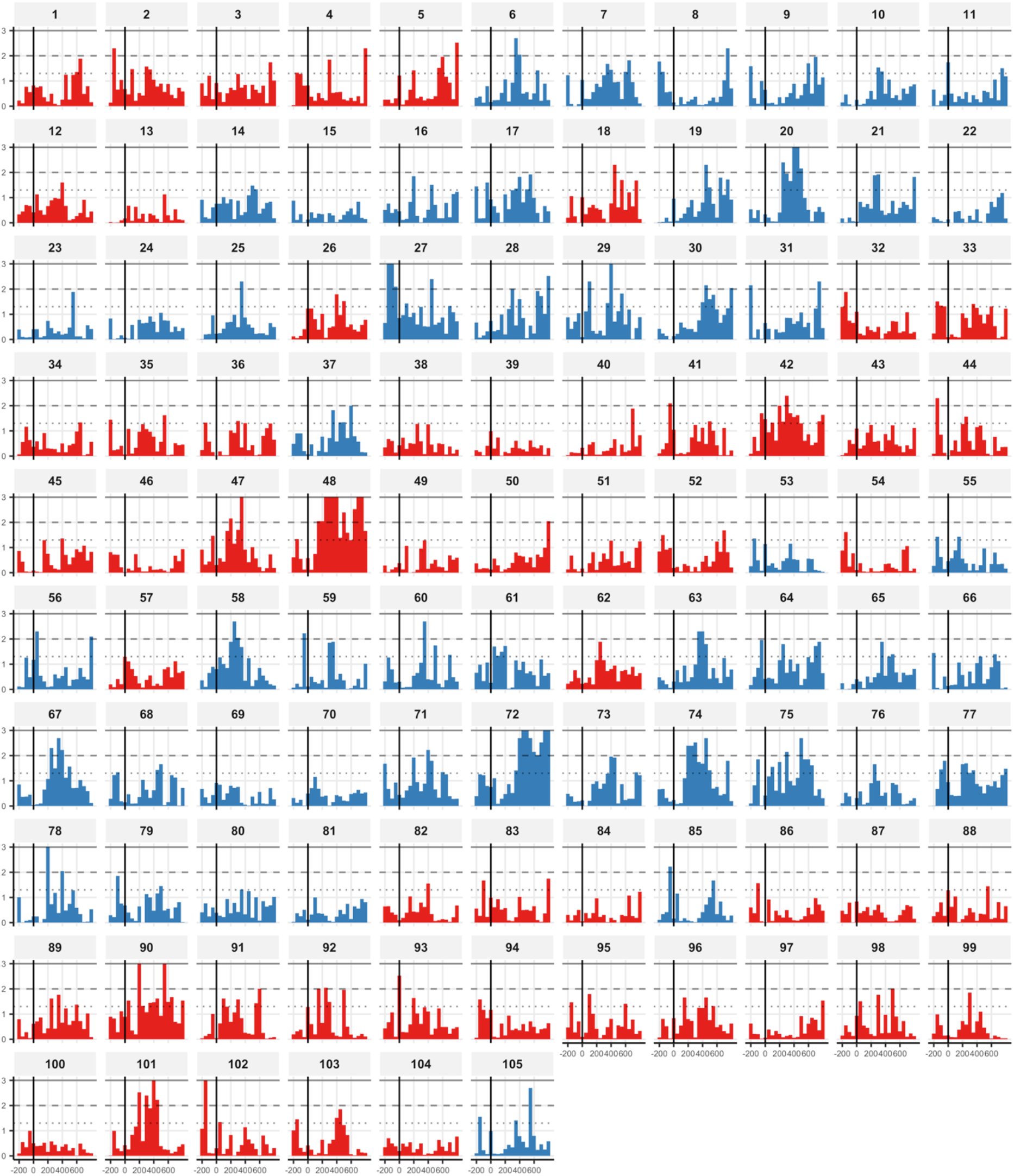
Participant-wise −log_10_ *p* value time series of the time-frequency-resolved EEG decoding in the theta frequency band. The statistical significance of within-participant EEG decoding per time bin was assessed using 1,000 permutations. The x-axis represents time (in ms) relative to stimulus onset; the y-axis represents −log_10_ *p* values. The solid black vertical line denotes stimulus onset. The dotted horizontal line at y ≈ 1.3 denotes *p* = 0.05, the dashed horizontal line at y = 2 denotes *p* = 0.01, and the solid horizontal line at y = 3 denotes *p* = 0.001. Absolute pitch musicians are colored in red; relative pitch musicians are colored in blue.

**Supplementary Figure 3A.**
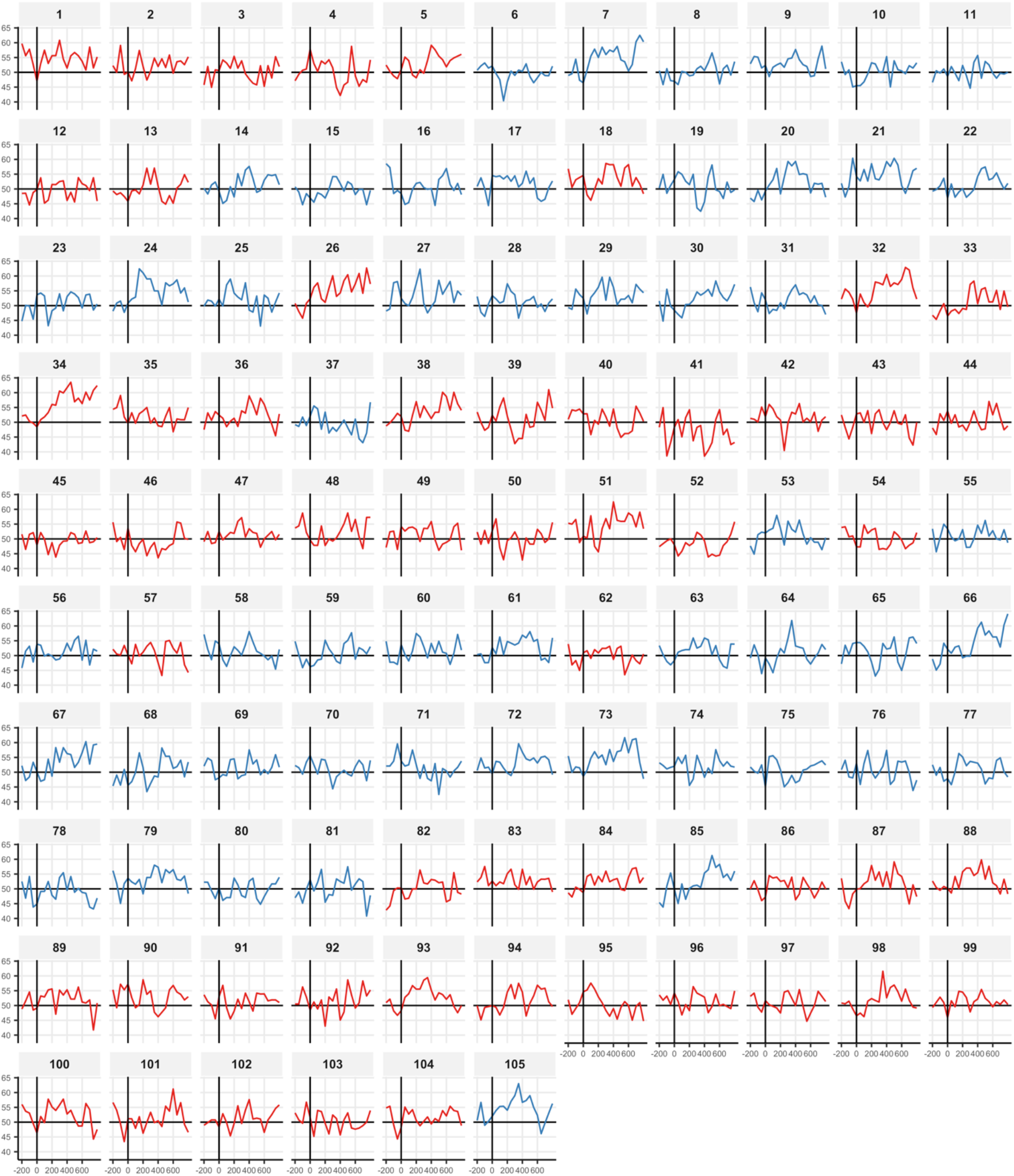
Participant-wise decoding time series of the time-frequency-resolved EEG decoding in the beta frequency band. The x-axis represents time (in ms) relative to stimulus onset; the y-axis represents decoding accuracy (in %). The solid black vertical line denotes stimulus onset. The solid black horizontal line at y = 50 denotes chance level (50%). Absolute pitch musicians are colored in red; relative pitch musicians are colored in blue.

**Supplementary Figure 3B.**
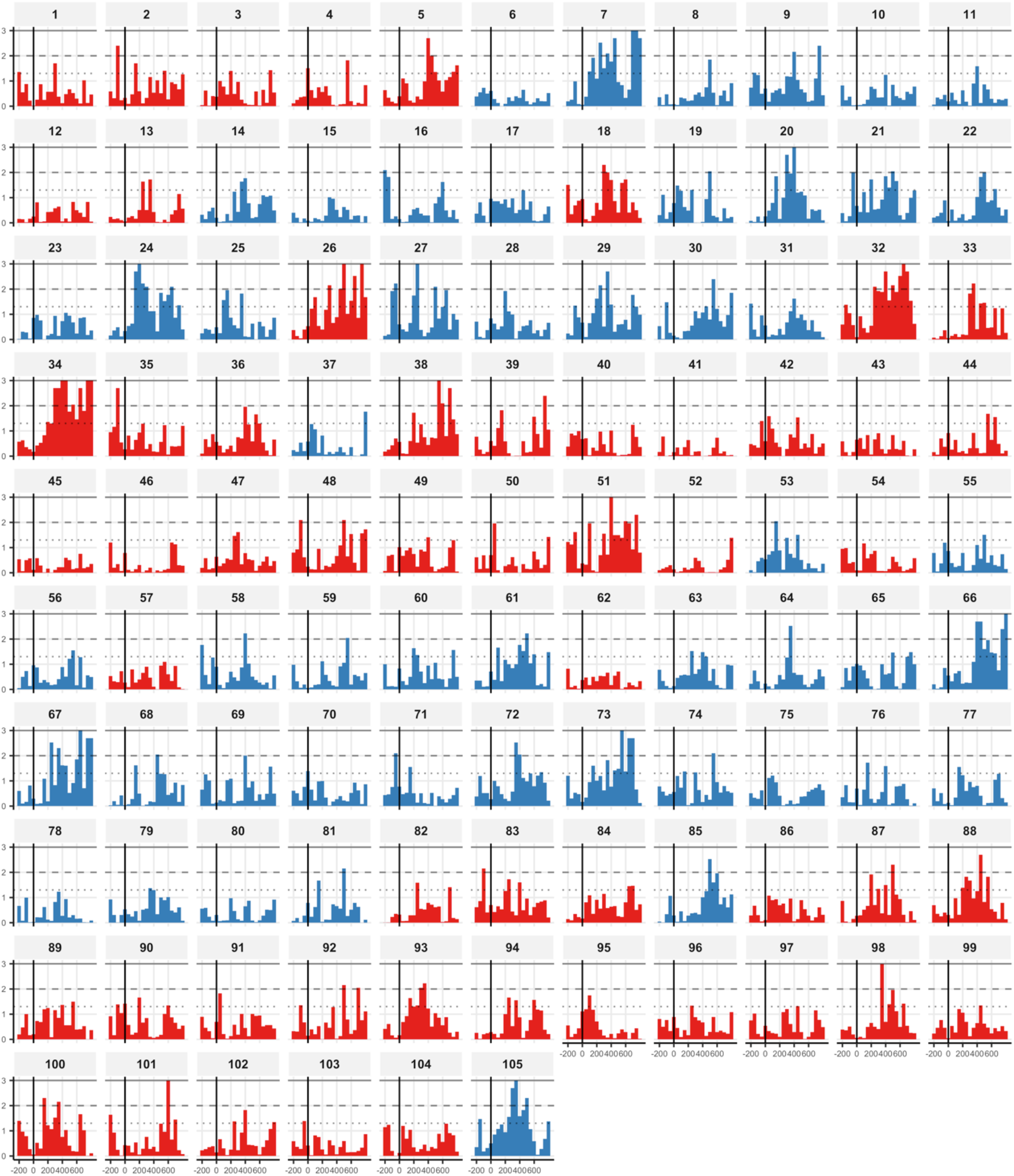
Participant-wise −log_10_ p value time series of the time-frequency-resolved EEG decoding in the beta frequency band. The statistical significance of within-participant EEG decoding per time bin was assessed using 1,000 permutations. The x-axis represents time (in ms) relative to stimulus onset; the y-axis represents −log_10_ *p* values. The solid black vertical line denotes stimulus onset. The dotted horizontal line at y ≈ 1.3 denotes *p* = 0.05, the dashed horizontal line at y = 2 denotes *p* = 0.01, and the solid horizontal line at y = 3 denotes *p* = 0.001. Absolute pitch musicians are colored in red; relative pitch musicians are colored in blue.

## Notes

https://dx.doi.org/10.17605/OSF.IO/7QXJS

